# Differential chromatin accessibility in developing projection neurons is correlated with transcriptional regulation of cell fate

**DOI:** 10.1101/645572

**Authors:** Whitney E. Heavner, Shaoyi Ji, James H. Notwell, Ethan S. Dyer, Alex M. Tseng, Johannes Birgmeier, Boyoung Yoo, Gill Bejerano, Susan K. McConnell

## Abstract

We are only just beginning to catalog the vast diversity of cell types in the cerebral cortex. Such categorization is a first step toward understanding how diversification relates to function. All cortical projection neurons arise from a uniform pool of progenitor cells that lines the ventricles of the forebrain. It is still unclear how these progenitor cells generate the more than fifty unique types of mature cortical projection neurons defined by their distinct gene expression profiles. Here we compare gene expression and chromatin accessibility of two subclasses of projection neurons with divergent morphological and functional features as they develop in the mouse brain between embryonic day 13 and postnatal day 5 in order to identify transcriptional networks that diversity neuron cell fate. We find groups of transcription factors whose expression is correlated with chromatin accessibility, transcription factor binding motifs, and lncRNAs that define each subclass and validate the function of a family of novel candidate genes *in vitro*. Our multidimensional approach reveals that subclass-specific chromatin accessibility is significantly correlated with gene expression, providing a resource for generating new specific genetic drivers and revealing regions of the genome that are particularly susceptible to harmful genetic mutations by virtue of their correlation with important developmental genes.

## Introduction

The cerebral cortex is the region of the human brain responsible for perception, language, complex thinking, and motor control. Neurons in the cortex can be subdivided into two broad classes: excitatory glutamatergic projection neurons and inhibitory GABAergic interneurons. Subclasses of cells exist within each of these broad classes, and recent efforts have sought to identify the complete catalog of cell types in the cortex using transcriptional profiling^1–4^. These studies have found that excitatory neurons are more diverse than inhibitory neurons, and at least 13 unique types of excitatory neurons have been identified in the developing mouse cortex^2^, while the adult mouse cortex contains at least 52^1,3^.

While our ability to identify neuron subtypes contributes to our understanding of functional divisions within cortical circuits^5^, how these circuits are specified during development remains unclear. The cerebral cortex is populated by young neurons in an inside-first, outside-last sequence: young excitatory projection neurons migrate from the subventricular zone and settle in discrete layers within the cortex. The deep layers (DL), layer 6 then layer 5, are populated first, followed in order by the upper layers (UL), layer 4 followed by layer 2/3^6^. Although DL and UL subclasses contain heterogeneous cell populations, cells within each subclass share common morphological and electrophysiological properties. The majority of DL neurons are corticofugal projection neurons (CFPNs), which send their axons to areas outside the cortex, including the thalamus and the spinal cord. The majority of UL neurons are cortico-cortical projection neurons (CPNs), many of which send their axons across the midline along the corpus callosum. While CPNs overwhelmingly occupy the upper layers, they can also be found among DL neurons.

A network of key transcription factors (TFs) expressed in early projection neurons determines whether a cell will acquire CFPN or CPN characteristics^7–9^. Previous studies in the mouse have revealed that TBR1 and FEZF2 regulate L6 and L5 CFPN development, respectively^10–17^, while SATB2 is required for CPN fate^18–20^. The downstream transcription cascade governing layer-specific maturation over time, however, is still unknown. No single study has directly assessed chromatin accessibility -- an important indicator of gene regulation -- *and* gene expression in CPNs or CFPNs during a specific stage of embryonic or early postnatal development. Moreover, it is still unknown how such transcriptional profiles would differ between CPNs and CFPNs at comparable stages of development. Previous screens have used RNA sequencing to identify genes that are significantly differentially expressed between projection neuron subtypes on the *same* day of development in the mouse, when DL neurons are more developmentally mature than UL neurons (Fig. 1A)^21^, or have relied on clustering algorithms and machine learning to define subtypes among groups of single cells^2,22^.

**Figure 1.**
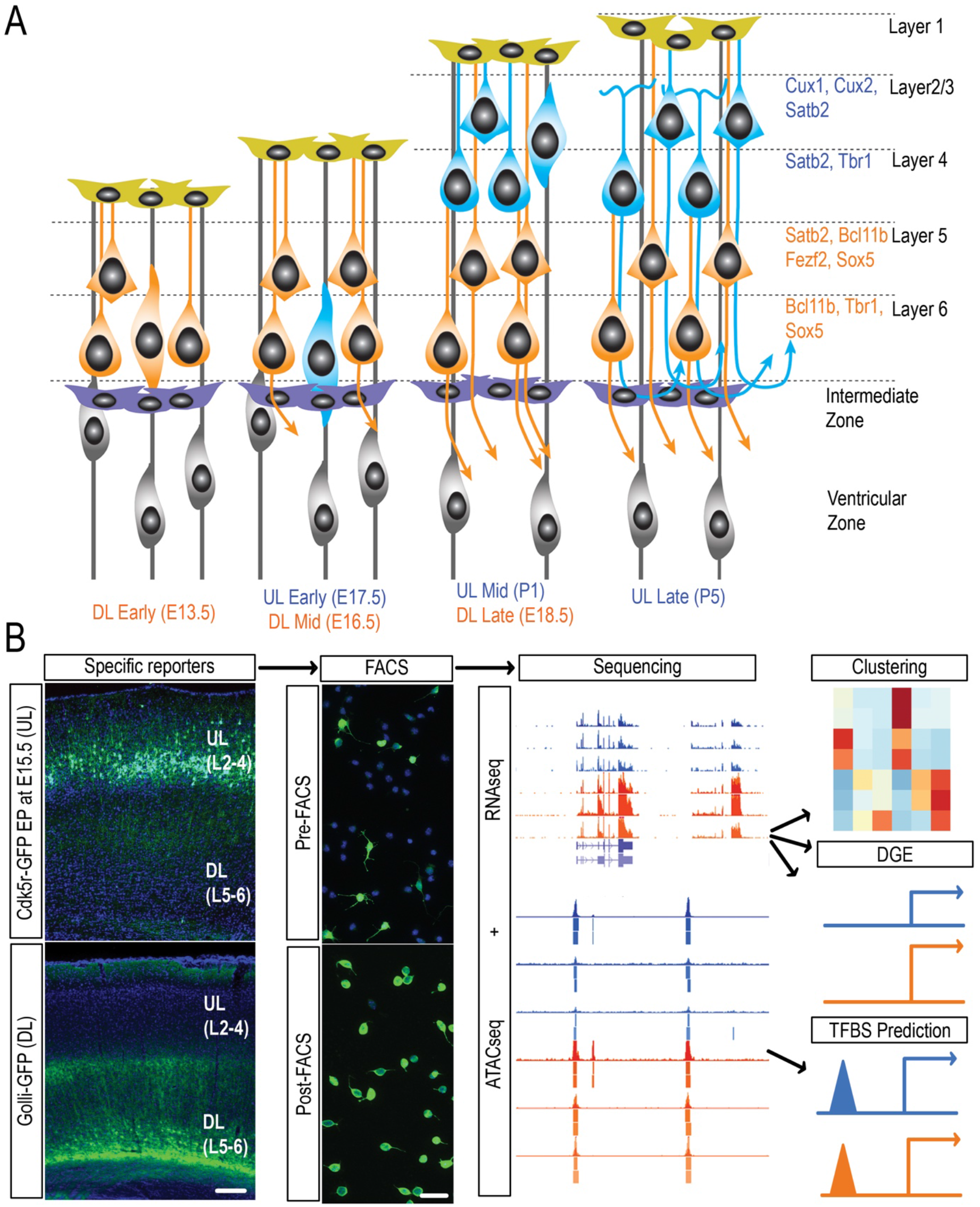
Excitatory projection neurons of the mouse neocortex can be subdivided into two broad classes, here called upper layer (UL) and deep layer (DL), each identified by a specific combination of transcription factors and labeled by an exclusive fluorescent reporter. **(A)** Schematic representation of neocortical development illustrating the stereotypical “inside-first, outside-last” sequence of cell fate specification and differentiation: DL neurons are generated first and reach milestones of development prior to UL neurons. **(B)** Experimental pipeline showing *in vivo* expression of cell class-specific reporters, FACS purification, and computational analysis. *(Bottom left)* Mouse cortex at P25 expressing Golli-GFP in DL neurons, L6 apical dendrites terminating in L4. *(Top left)* mouse cortex at P9 expressing Cdk5r-GFP in UL neurons occupying L2/3. Scale bars are 100 μm for *in vivo* expression and 20 μm for dissociated cells.

Here, we analyze cell type-specific gene expression and differential chromatin accessibility over three defined stages of development in two sets of genetically identified neurons, and we compare our results with the results of previous sequencing screens in mouse and human cortex. We find broad agreement between our bulk-sequenced neuron populations and populations identified by single cell sequencing. Moreover, given the robustness of bulk sequencing, we are able to join open chromatin with gene expression within these defined populations and predict transcription factors with previously uncharacterized roles in specifying UL versus DL fate. In addition, we identify long non-coding RNAs adjacent to important TF genes (“TF-lncRNAs”) with highly specific spatial and temporal expression patterns. Finally, we test the ability of select candidate genes to modulate the ratio of CPN to CFPN neurons *in vitro*.

## Results

### Cell subclass is the greatest source of variability in gene expression

Deep layer projection neurons (DL subclass, layers 5 and 6) are generated first and mature more quickly than upper layer projection neurons (UL subclass, layers 2-4) (Fig. 1A). Here, we define three stages of maturation common to DL and UL neurons during the initial period after exiting the cell cycle: 1) “early,” when newly post-mitotic cells are entering or have just entered the cortical plate; 2) “mid,” during primary axon extension; and 3) “late,” during axon collateral formation^23–25^.

We established a protocol for isolating DL and UL neurons at these three stages (Fig. 1B). For DL neurons, we used fluorescent-activated cell sorting (FACS) of micro-dissected and dissociated mouse neocortex expressing the transgene *golli*-τ-EGFP (*golli*-EGFP), in which the 1.3 kb *golli* promoter of the myelin basic protein drives expression of EGFP in the subplate and in corticothalamic, corticospinal, corticocallosal, and corticocollicular neurons of layers 5 and 6^26^. FACS purification of GFP-positive cells was performed on embryonic day (E) 13.5 (Theiler Stage 21) (“DL early”), E16.5 (TS24) (“DL mid”), and E18.5 (TS 26) (“DL late”). For UL neurons, we introduced a birth-dating plasmid containing the cyclin-dependent kinase 5 regulatory subunit p35 (Cdk5r) promoter driving GFP by *in utero* electroporation at E15.5, during the peak of UL neurogenesis. The Cdk5r promoter is expressed in neurons after cell cycle exit^27–29^. We were therefore able to use GFP expression to FACS-purify UL cortico-cortical projection neurons, primarily consisting of layer 2/3 corticocallosal cells, at E17.5 (TS25) (“UL early”), postnatal day (P) 1 (“UL mid”), and P5 (“UL late”).

We used RNAseq to quantify gene expression at early, mid, and late stages of DL and UL development. We used two biological replicates per stage per subclass and generated an average of 2.35×10^7^ reads per replicate (range, 1.65×10^7^ − 2.86×10^7^), each replicate having an adequate number of reads with which to compare gene expression across space and time (Fig. S1A). These data have been deposited in GEO and can be accessed under the accession number GSE116147.

Previous reports have suggested that in the developing neocortex, cell subclass does not contribute as much variability to gene expression as does developmental time^21^. Recent single-cell sequencing data, however, have demonstrated that gene expression can distinguish cell subclass^1–3,30^. To test whether cell subclass is the greatest source of variability in gene expression, we used PCA, t-SNE, or hierarchical clustering of DL and UL gene expression over time to identify clusters of cell types in the combined dataset. PCA was conducted on all replicates at every condition for a total of 12 points. For each replicate, PCA first used all Ensembl mm9 transcripts with UCSC mappings (34,307 transcripts) as features and showed that the first two principal components were able to separate UL from DL replicates; however, two of the DL replicates clustered closely with the UL replicates. Two-dimensional t-distributed stochastic neighbor embedding (t-SNE) using all transcripts showed even clearer clustering of all replicates within a subclass. (Fig. S1B). Due to our interest in transcription cascades, when we limited the PCA analysis to transcripts encoding transcription factors (1,461 transcripts), replicates of the same stage and cell subclass clustered more closely (Fig. 2A). These data suggest that gene expression, particularly that of TFs, is able to distinguish cell subclass.

**Figure 2.**
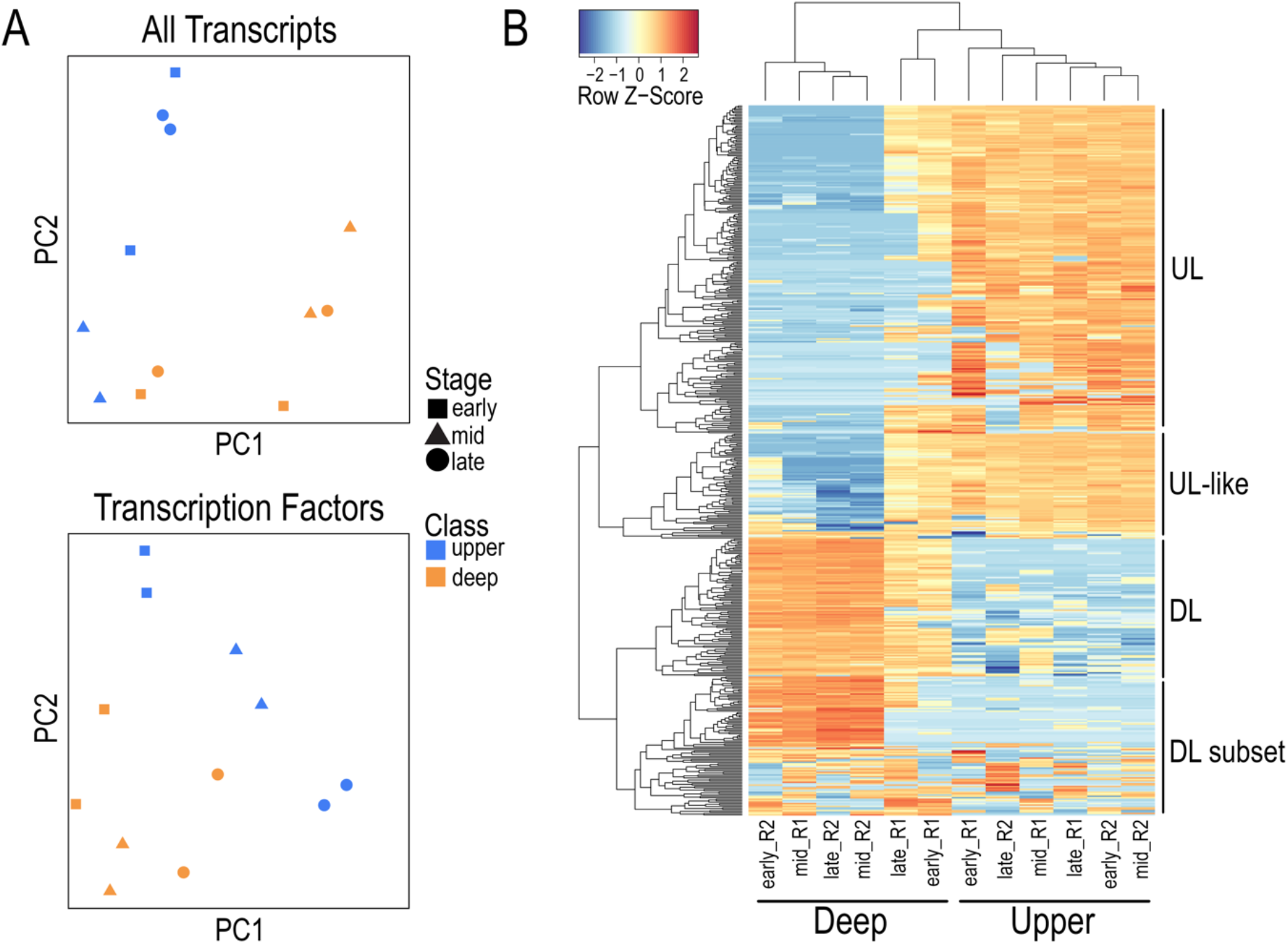
Principal component analysis (PCA) and naive hierarchical clustering show that gene expression is able to distinguish cell subclass. **(A)** PCA of all transcripts and of all transcription factor transcripts shows that variance between cell subclass is captured in the first two principal components. **(B)** Naive hierarchical clustering of the top 500 most variable Ensembl mm9 transcripts separates UL from DL neurons and identifies a cluster of “UL-like” genes in DL neurons. Clusters labeled to the right are listed in Table S1.

On the 500 transcripts with the most variability in gene expression across both subclasses, hierarchical clustering revealed that two of the six DL replicates (one early and one late) clustered more closely with the UL replicates than with the other four DL replicates (Fig. 2B). Given that the deep layers of the neocortex consist of *both* cortico-cortical projection neurons (CPNs) and corticofugal projection neurons (CFPNs), while upper layers primarily consist of CPNs alone, we hypothesized that a sub-cluster of highly variable transcripts defines a population of UL-like cells that populate both deep and superficial layers. We therefore identified four sub-clusters of transcripts that breakdown our two subclasses into four putative sub-subclasses: “UL” transcript cluster defining UL neurons, “UL-like” transcript cluster defining some UL-like DL neurons, “DL” transcript cluster defining DL neurons, and “DL subset” transcript cluster defining a subset of DL neurons. Each cluster contains known markers of UL or DL identity (***Supplemental Table 1***): *Nfix* and *Cux1* in the *“*UL” cluster; *Sox5* and *Tle4* in the “DL” cluster; *Ptn*, a gene expressed in DL CPNs in both mouse and macaque, in the “UL-like” cluster^31^, and the LIM-domain containing gene *Lmo3*, which is expressed in DL neurons, in the “DL-subset” cluster^32^. The three genes that encode the subunits of the platelet-activating factor acetylhydrolase 1B complex (PAFAH1B2), an enzyme involved in neuron migration and synaptic function, were also differentially clustered: *Pafah1b1* (*Lis1*) and *Pafah1b2* clustered with DL transcripts, and *Pafah1b3* clustered with UL transcripts, consistent with a previous report showing different expression patterns of these subunits^33^. Moreover, when we limited hierarchical clustering to only those transcripts that encode TFs, UL and DL subclasses clearly segregated (Fig. S1C).

In order to better characterize these four sub-clusters, we compared them with the cortical cell types identified using single-cell sequencing of brain cells from P2 and P11 mouse (“SPLiT-seq cells”)^2^ (Table 1). DL and DL-subset cells shared expression with the DL SPLiT-seq cell types, while UL cells shared expression with UL SPLiT-seq cell types. UL-like cells expressed *Mpped1*, a gene expressed in both UL and DL SPLiT-seq cell types. Likewise, when we compared gene expression of our sub-clusters with genes expressed during UL (L4) differentiation^34^, only UL sub-cluster genes significantly overlapped with L4 genes (hypergeometric *P*-value: 1.0×10^−4^ for UL genes/L4 “wave 2” genes and 1.3×10^−7^ for UL genes/L4 “wave 5” genes). DL genes and L4 genes did not significantly overlap, after correcting for multiple comparisons. Together, these data demonstrate that our bulk populations represent transcriptionally distinct UL and DL projection neuron subclasses.

**Table 1.** Shared expression of subclass-defining genes identified by bulk RNAseq or single cell RNAseq

| Sub-class | L2/3/4-Mef2c | L2/3/4-Ntf3 | L4-Rorb | L4-Wnt5b | L4/5 | L5/6-Npr3 | L5/6-Sulf1 | L5-Fezf2 | L6 | L6a |
| --- | --- | --- | --- | --- | --- | --- | --- | --- | --- | --- |
| UL | *Cux1* |  | *Cux1* | *Meis2*,<br>*Nfix*,<br>*Ppfia2* |  |  |  |  |  |  |
| UL-like | *Mpped1* | *Mpped1* |  | *Mpped1* |  | *Mpped1* |  |  | *Mpped1* | *Mpped1* |
| DL |  |  |  |  |  | *Sox5*, *Tle4* | *Tle4* | *Tle4* | *Sox5*,<br>*Tle4* | *Cdh13*,<br>*Sox5* |
| DL-subset |  |  |  |  | *Lmo3* | *Lmo3*,<br>*Dlg2* |  |  |  |  |

### DL and UL neuron subclasses can be defined by consistently significant differential expression of TFs over time

Given that gene expression, particularly that of TFs, can distinguish DL from UL identity, we next asked which transcripts are significantly different between DL and UL neurons at all three stages of development observed, and whether such transcripts are differentially enriched for known biological functions.

We first identified all transcripts that are significantly higher in DL or UL neurons at all three stages (*Q-*value <= 0.05 for each pairwise comparison) (***Supplemental Table 2***). The 199 transcripts that were consistently higher in DL neurons were enriched for genes involved in neuron commitment in the forebrain (GO:0021902) and mRNA splice site selection (GO:0006376), while the 235 transcripts that were consistently higher in UL neurons were enriched for genes involved in glucocorticoid receptor signaling (GO:2000324) and postsynaptic membrane organization (GO:1901628). Within these two groups of transcripts, we identified 26 TFs that are consistently higher in DL neurons, including the known markers *Tbr1* and *Nfib*, and 21 TFs that are consistently higher in UL neurons, including the known marker *Pou3f2* (Fig. 3A,B, ***Supplemental Table 2***). DL neurons also expressed the recently identified DL-specific CPN marker *Rprm*, and UL neurons expressed the UL-specific CPN marker *TNC*^35^. Potentially novel subclass markers with previously characterized roles in neural development and function included *Zfp521* (DL) and *Bhlhe40 (Bhlhb2/Dec1/Sharp1*) (UL) (Fig. 3D)^36,37^.

**Figure 3.**
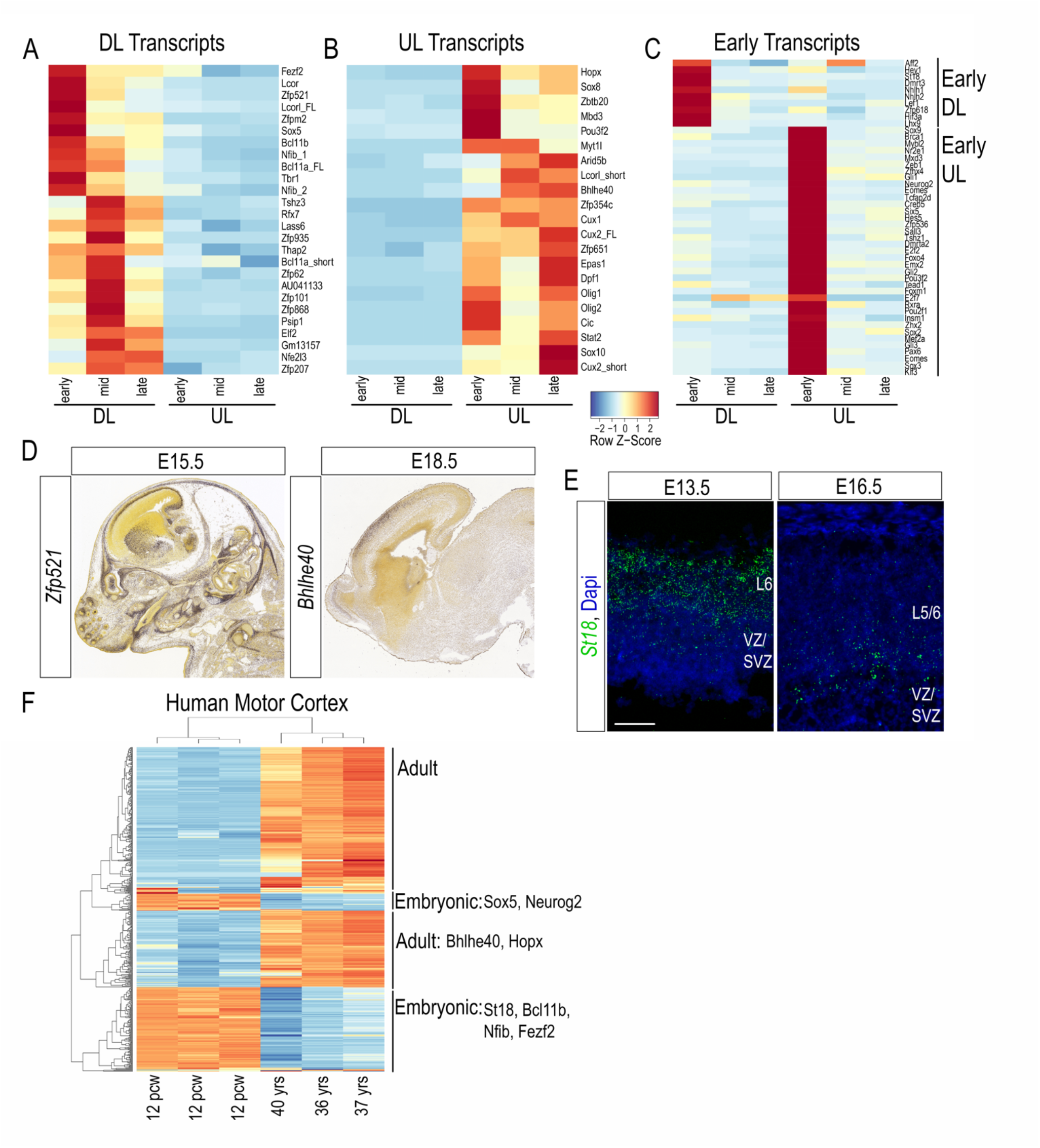
Specific expression of a set of TFs throughout development distinguishes UL from DL populations. **(A, B)** Heatmaps of relative expression of TFs that are significantly higher in DL (A) or UL (B) neurons at all three stages of development. FL stands for full length. **(C)** Heatmap of relative expression of DL- and UL-specific “early” TFs. For all three heatmaps, each column represents the average transcripts per million (TPM) of two replicates. **(D)** *In situ* hybridization of the DL TF *Zfp521* and the UL TF *Bhlhe40* on sections through the embryonic mouse brain showing restricted expression in the cortex. *Image Credit: Allen Institute*. **(E)** Fluorescent *in situ* hybridization of the DL early TF *St18* on sections through the embryonic mouse cortex showing enriched expression in layer 6 at E13.5. **(F)** Naïve hierarchical clustering of the top 500 most variable transcripts between embryonic and adult human motor cortex reveals expression of DL and UL TFs in the embryonic cluster and two UL TFs in the adult cluster. Scale bar in E is 50 μm.

To identify subclass-specific TFs relevant to human cortical development, we compared our data with RNAseq data from embryonic (12 pcw) and adult (36-40 years) human motor cortex from the BrainSpan atlas^38^. Naive hierarchical clustering of the top 500 most variable BrainSpan transcripts showed clear segregation of embryonic and adult motor cortex gene expression (Fig. 3F). Six mouse subclass-specific TFs were enriched in the embryonic human motor cortex clusters. Surprisingly, two mouse UL TFs, *Bhlhe40* and *Hopx*, were enriched in the *adult* human motor cortex clusters, suggesting that early embryonic TFs identified in mouse UL neurons are relevant to human motor cortex development and function.

Given its UL-specificity and relevance to human motor cortex, we searched for potential protein binding partners of Bhlhe40 using the STRING database^39^. A regulator of circadian rhythm, Bhlhe40 is known to interact with the clock genes *Clock*, *Cry1*, *Npas2*, *Nr1d2*, and *Arntl* (Fig. S2A). Of these, *Arntl* also shows significantly higher expression in UL neurons (Fig. S2B, ***Supplemental Table 8***). Moreover, *Arntl* and *Bhlhe40* both interact with the UL-enriched TF gene *Epas1* (***Supplemental Table 2***), which is expressed in UL neurons at P4, similar to *Bhlhe40* (Fig. S2C,D), and opposite the layer 5-enriched *Bcl11b* (Fig. S2E).

### Olig2-expressing cells are generated from late progenitor cells

We were surprised to observe *Olig1* and *Olig2* expression in Cdk5r-GFP+ neurons, given that the Cdk5r promoter has been reported to be specific to excitatory neurons in the cortex, and cortical expression of *Olig1* and *Olig2* is specific to oligodendrocyte progenitor cells (OPCs)^40,41^. A previous observation, however, suggests that at least a subset of OPCs express Cdk5r^42^. RNAseq read pile-ups that map to the *Olig2* locus show enriched expression in the 3’UTR in the mid and late stages, compared with the early stage, suggesting that cells fated to become UL neurons may maintain low expression of *Olig2* mRNA early after cell cycle exit and retain expression of the 3’UTR later in development, as has been demonstrated for other transcripts in the nervous system (Fig. S3A)^43^.

In order to determine whether a subset of Cdk5r-GFP+ cells are in fact OPCs, we quantified the percentage of Olig2+ cells in the Cdk5r-GFP population. We used immunohistochemistry to co-label GFP-electroporated cortices with an antibody to Olig2 at E17.5, P3, and P9 and found that a small subset of Cdk5r-GFP+ cells maintained expression of Olig2 as late as P9 (Fig. S3C-F). The percentage of co-labeled cells, however, ranged from only 1.2%-4.0% per embryo and thus represented only a small fraction of total Cdk5r-GFP+ cells (Fig. S3B), agreeing with a previous study showing that the progeny of cortical progenitor cells at E14.5 contains a percentage of oligodendrocytes^44^.

### Subclasses of projection neurons have common and specific expression dynamics

Common and specific gene expression dynamics may reveal shared and distinct biological functions of these projection neuron subclasses. To identify groups of genes that are specific to a cell subclass at one stage of development and genes that are common to both cell subclasses at a specific stage of development, we compared gene expression across time then space to arrive at sets of genes that define these two dimensions of development.

We strictly defined a transcript specific to the early stage as being significantly higher in the early stage compared with both the mid stage AND the late stage of the same cell subclass. TFs that were found to be “early” in only one cell subclass were highly restricted to the early stage in that cell subclass (Fig. 3C). We used fluorescent *in situ* hybridization (FISH) to confirm early DL expression of *St18*, a member of the Myt family of TFs. *St18* was highly expressed in layer 6 at E13.5, and its expression was reduced dramatically in the cortical plate by E16.5 (Fig. 3E).

We next asked if there were “early,” “mid,” or “late” transcripts that are common between the two cell subclasses. Here we defined “early” and “late” as being significantly higher (*Q-value* <= 0.05, log2FC >= 1.5) in the early stage or late stage compared with the middle stage, regardless of expression at the other end. Likewise, we defined a “mid” transcript as being significantly higher in the middle stage compared with the early stage but not with the late stage. There were many more common “early” TFs than common “mid” or “late” TFs, and many of these were highly restricted to the early stage in both cell subclasses (Fig. S4A, ***Supplemental table 3***). We also identified additional “mid” and “late” TFs specific to one cell subclass (Fig. S4B, ***Supplemental table 4***) and axon guidance genes specific to each stage but not subclass-specific (Fig. S4C).

A recent study used single cell RNA sequencing (scRNAseq) of embryonic mouse brain to identify cohorts of genes that define stages of neuron development across time, naïve to cell type, and concluded that early gene expression is enriched for cell-intrinsic transcriptional regulators, while later-enriched genes suggest a shift to cell-extrinsic mechanisms^22^. Similarly, Gene Ontology (GO) analysis^45^ of our common stage-specific transcripts showed a shift from transcriptional regulation by the Smoothened pathway, which was enriched among “early” genes, to cell-extrinsic signaling across the synaptic cleft for “mid” genes, and positive regulation of cell communication (interactions between a cell and its surroundings) for “late” genes (Fig. S4C,D). Moreover, there was significant overlap of our early gene set and genes that define late “basal progenitors” among sequenced single cells (hypergeometric *P*-value: 3.50×10^−18^). Likewise, our mid gene set significantly overlapped genes that define single DL neurons four days after differentiation (hypergeometic *P*-value: 3.25×10^−14^) as well as genes that define single UL neurons four days after differentiation (hypergeometric *P*-value: 1.42×10^−19^). Our late gene set was beyond the latest time point analyzed by scRNAseq and therefore did not significantly overlap any identified gene cluster. These data suggest agreement between our bulk-sequenced populations and populations identified by scRNAseq.

### TF-lncRNAs have cell-specific expression dynamics

Highly specific expression of a transcript at a single stage of development may indicate a specific developmental function. In order to identify genes that are highly specific at a given stage, we looked for transcripts that “spike” at the mid stage of development, or show high expression at the “mid” stage relative to “early” and “late,” (*q-value* <= 0.05, log2FC >= 1.43) and found only a few transcripts that spike in either DL neurons or UL neurons (***Supplemental Table 5***). Surprisingly, three of the five DL “spike” transcripts are long non-coding RNAs (lncRNAs) that are each adjacent to a major cortical TF gene. We here call these transcripts *Satb2-lncRNA (9130024F11Rik)*, *Pou3f3-lncRNA* (2900092D14Rik), and *Pou3f2-lncRNA* (AK039117), because they are adjacent to *Satb2*, *Pou3f3* (*Brn1*), and *Pou3f2* (*Brn2*), respectively. In DL neurons, *Satb2-lncRNA* and *Pou3f3-lncRNA* parallel the expression dynamic of their neighboring TF gene, supporting previous observations that lncRNA expression is often correlated with that of a proximal gene with which it shares a bidirectional promoter^46–49^. Of these three lncRNA transcripts, however, only *Satb2-lncRNA* showed a similar, though minor spike in expression in UL neurons (*Q-value: 0.01* and *0.12* for early vs. mid and mid vs. late, respectively), mirroring the expression dynamic of *Satb2* (Fig. 4A).

**Figure 4.**
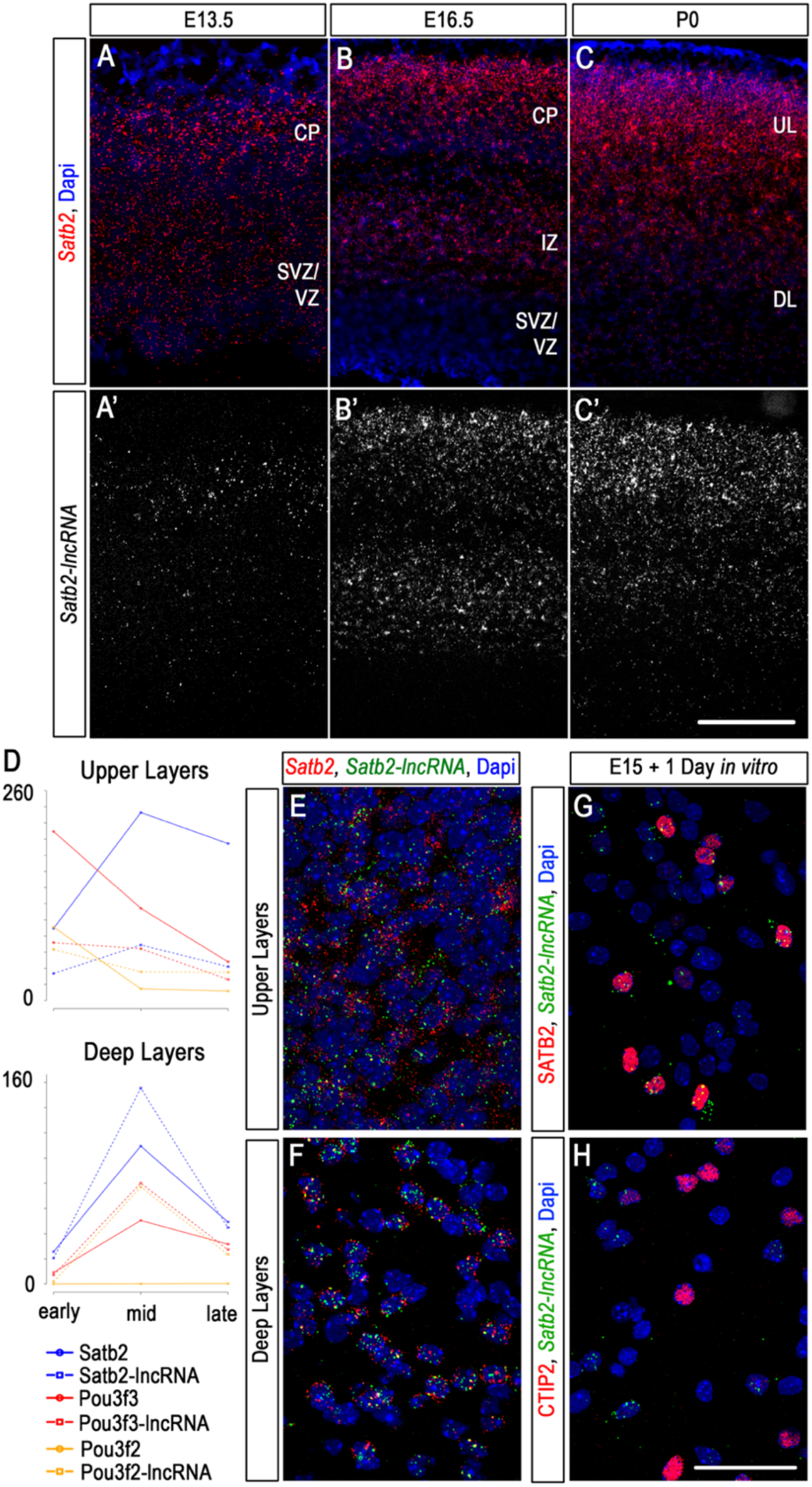
*Satb2-lncRNA* is differentially expressed in cortical neurons over time and space. **(A-C’)** Fluorescent *in situ* hybridization (FISH) of *Satb2* (A-C) and *Satb2-lncRNA* (A’-C’) in the mouse neocortex at E13.5, E16.5, and P0. **(D)** TF-lncRNAs show increased expression (in FPKM) during mid stage in DL neurons, and *Satb2-lncRNA* shows a similar spike in UL neurons. **(E-H)** FISH of *Satb2-lncRNA* co-labeled with *Satb2* RNA (E,F), SATB2 protein (G), or CTIP2 protein (H) showing expression in UL nuclei and cell bodies (E), DL nuclei (F), and SATB2-positive cells and exclusion from CTIP2-positive cells (H). Scale bar in C’ is 200 μm for A-C’. Scale bar in H is 50 μm for E-H.

Two additional lncRNAs that flank *Pou3f3* have been shown to regulate cortical progenitor proliferation and cell fate in the mouse^50,51^. We hypothesized that *Satb2-lncRNA* may play a similar role in cell fate. We used two-color FISH to analyze the localization of *Satb2-lncRNA* relative to that of *Satb2* (Fig. 4B-F). We found that *Satb2-lncRNA* expression parallels that of *Satb2* in the developing neocortex. Both transcripts are expressed in the cortical plate (CP) and in the ventricular zone (VZ) at E13.5, becoming restricted to the CP and intermediate zone (IZ) by E16.5 and enriched in UL neurons by P0.

Confocal imaging confirmed that individual cells co-express both *Satb2* and *Satb2-lncRNA* transcripts; however, their sub-cellular localization differs between DL neurons and UL neurons. In DL neurons, their expression is restricted to the nucleus, whereas in UL neurons, their expression appears both nuclear and cytoplasmic (Fig. 4E,F). Moreover, *Satb2-lncRNA* was expressed in cells positive for Satb2 protein but not in cells positive for Ctip2 protein (Fig. 4G,H).

The co-expression of *Satb2* and *Satb2-lncRNA* and the differential sub-cellular localization in DL vs. UL neurons suggest that *Satb2-lncRNA* may play a role in regulating the balance of SATB2+ CPNs and CTIP2+ CFPNs in the neocortex. To test the hypothesis that *Satb2-lncRNA* is involved in a network regulating the choice between CPN and CFPN, we designed two siRNA constructs targeting the 3’ end of the first exon of *Satb2-lncRNA*. The first exon is shared by the short and long *Satb2-lncRNA* isoforms and does not overlap the first exon of *Satb2* (Fig. S5A). We confirmed efficient knockdown (KD) of expression of both isoforms of *Satb2-lncRNA* and diminished cytoplasmic *Satb2-lncRNA* localization in dissociated cortical neurons cultured for 72 hours (Fig. S5B,C). To evaluate the effect of KD on cell fate, we co-labeled control (scrambled siRNA) and KD (siRNA 1) treated cells with antibodies to SATB2 and CTIP2 and counted the ratio of SATB2+ to CTIP2+ cells (Fig. S5D,E). The number of cells expressing SATB2 relative to CTIP2 was reduced modestly but not significantly (*P* = 0.055) in KD cells compared with control cells. Together, these results suggest that *Satb2-lncRNA* may be involved in regulating the balance of CPNs to CFPNs; however, cytoplasmic-specific reduction of *Satb2-lncRNA* had a negligible effect on cell fate.

### ATACseq of subclasses of projection neurons suggests specific TFs regulate subclass-specific gene expression

To identify additional pathways involved in regulating the balance of CPNs to CFPNs, we used an assay for transposase accessible chromatin using sequencing (ATACseq) to compare chromatin accessibility between DL neurons and UL neurons at early, mid, and late stages (Fig. 5A)^52^. ATACseq datasets generated here are available in GEO (accession number GSE116147).

**Figure 5.**
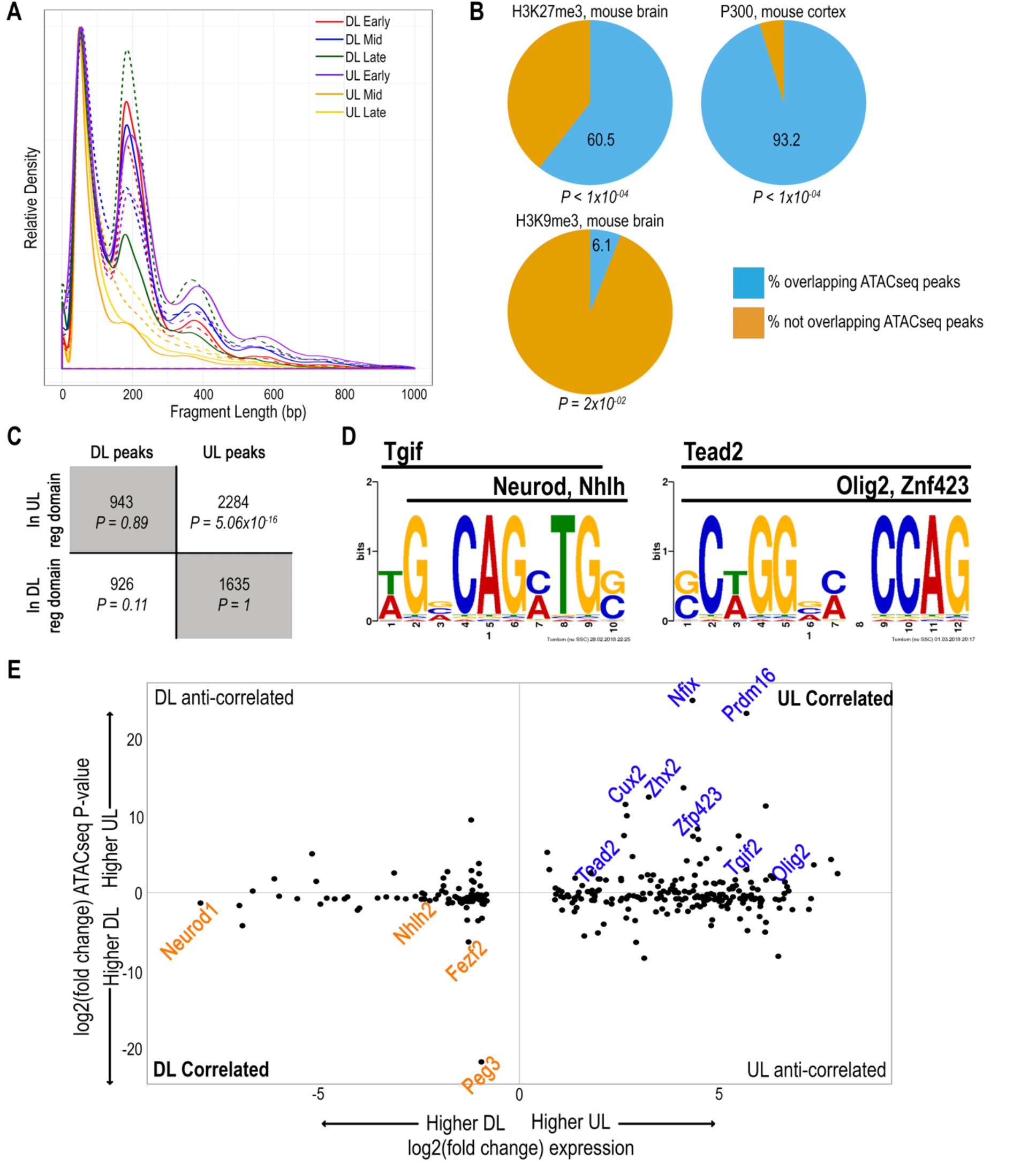
Chromatin accessibility is associated with gene expression. **(A)** The density of fragment lengths for each ATACseq library. Each stage in each cell class has two biological replicates, represented by a solid line and a dotted line. **(B)** The union of ATACseq peaks significantly overlaps previously identified cortical enhancers and repressors. **(C)** Binomial test showing that a class-specific ATACseq peak is more likely to occur in the regulatory domain of a gene more highly expressed in that same class. **(D)** Enriched motifs and predicted TF binding sites found in the union of ATACseq peaks of correlated DL genes (left) and UL genes (right). **(E)** Early TFs that are significantly differentially expressed between DL and UL neurons plotted by their log2(fold change) in expression and their log2(fold change) in binomial *P*-value for class-specific ATACseq peaks. Each comparison is UL:DL, so TFs with negative values for each are “correlated DL” TFs, and TFs with positive values for each are “correlated UL” TFs. TFs with the greatest change in binomial *P*-value or are referenced in Figures 5 and 6 are labeled.

TFs can bind DNA and modulate gene expression in regions where chromatin is accessible to transposase insertion, at putative promoters, enhancers, and repressors. Our complete set of early, mid, and late DL and UL ATACseq peaks was significantly enriched for previously identified enhancers and repressors in the mouse brain: 93.2% of cortical enhancers bound by P300 (*P*-value < 1×10^−4^), 60.5% of cortical repressors marked by H3K27me3 (*P*-value < 1×10^−4^), and 6.1% of cortical repressors marked by H3K9me3 (*P*-value < 2×10^−2^) overlapped our set of ATACseq peaks, confirming that these peaks represent both activating and repressing chromatin states (Fig. 5B).

After masking for blacklist regions, repeat regions, segmental duplications, and exons, we identified ATACseq peaks that were present in both replicates of each cell subclass at each stage (“replicate peaks”). The number of replicate peaks decreased between the early and mid/late stages for both DL and UL neurons (***Supplemental table 6***). Only a relative handful of replicate peaks in each subclass were both present in that subclass at all three stages *and* absent from the other subclass at all three stages. Moreover, within these sets, even fewer replicate peaks overlapped previously identified brain repressors (H2K27me3 ChIPseq) and enhancers (P300 ChIPseq)^53,54^ (Table 2). We hypothesized that these subclass-specific enhancers may represent regions important for establishing DL versus UL fate. We therefore used the GREAT regulatory domains to associate subclass-specific enhancers with nearby genes^55^ (***Supplemental table 7***). At least one of these genes has been shown to play a cell-type specific role in the neocortex: *Khdrbs1/Sam68* is known to regulate alternative splicing of *Nrxn1 and has an UL-specific enhancer just downstream of its 3’UTR*^56^.

**Table 2.** Peaks overlapping putative cortical enhancers and repressors

|  | **Number specific**<br>**peaks** | **Number overlapping**<br>**p300 ChIPseq** | **Number overlapping**<br>**H3K27me3 ChIPseq** |
| --- | --- | --- | --- |
| Upper Layers | 185 | 9 | 37 |
| Deep Layers | 1082 | 7 | 31 |

### The top conserved subclass-specific enhancers putatively regulate genes with subclass-specific expression

We next asked whether subclass-specific peaks are enriched near subclass-specific genes. An analysis of fragment length density revealed similar size distributions between early DL and early UL ATACseq sets, both of which had the highest numbers of replicate peaks (Fig. 5A). We therefore looked for subclass-specific regulatory elements in early DL and early UL neurons. We first searched for subclass-specific enhancers by identifying DL-specific and UL-specific ATACseq peaks overlapping cortical P300 ChIPseq peaks then ranking them by conservation (***see methods***). We identified 110 DL-specific enhancers and 1047 UL-specific enhancers. Of these, the top most conserved peaks were found to be in the regulatory domains of genes either known to be involved in cortical development or shown to have layer and/or stage-specific expression in the current study (Table 3).

**Table 3.** Top five conserved P300-bound enhancers in each subclass and known expression pattern of nearest gene

| Sub-class | Coordinates | Nearest Gene | Expression | Figure or citation |
| --- | --- | --- | --- | --- |
| UL | chr8:93432038-93432088 | Chd9 | Higher DL, mid stage |  |
| UL | chr14:118629169-118630069 | Sox21 | Early UL TF | Supp. table 4 |
| UL | chr3:5224759-5225409 | Zfhx4 | Early UL TF | Fig. 3C; Kostich and Sanes 1995 |
| UL | chr1:6724704-6725154 | St18 | Early DL TF | Fig. 3E and 6B |
| UL | chr7:122077180-122077730 | Gm6816 |  |  |
| DL | chr2:61636113-61636413 | Tbr1 | DL | Fig. 3A |
| DL | chr2:125327097-125327197 | Fbn1 | Common mid gene | Supp. table 3 |
| DL | chr3:100909214-100909314 | Ptgfrn | Higher UL, early and mid |  |
| DL | chr14:105894929-105894979 | 4930449E01Rik |  |  |
| DL | chr6:144145936-144146086 | Sox5 | DL | Fig. 3A |

These data suggest that subclass-specific chromatin accessibility may be correlated with cortical gene expression. We therefore hypothesized that subclass-specific ATACseq peaks are enriched next to subclass-specific genes, or genes that are significantly differentially expressed between DL and UL neurons. Using our RNAseq data from early UL and DL cells, we identified genes that were significantly differentially expressed (*Q*-value < 0.05), establishing a set of subclass-specific genes (1967 DL genes and 2706 UL genes). We then identified ATACseq peaks in the regulatory domains of subclass-specific genes, excluding all peaks that were common to both subclasses (1869 DL-specific peaks and 3919 UL-specific peaks) (Fig. 5C). Next, using a binomial *P*-value, which accounts for differences in regulatory domain sizes, we calculated the likelihood that a subclass-specific ATACseq peak would be in the regulatory domain of a gene that was specific to the same subclass, or “correlated” with gene expression. An “anti-correlated” pair, on the other hand, would consist of a subclass-specific peak localized to the regulatory domain of a subclass-specific gene in the opposing subclass. Of the 1869 peaks specific to the DL subclass, 926 localized to the regulatory domains of DL genes (binomial *P-value*: 0.11), while 943 localized to the regulatory domains of UL genes (binomial *P-value*: 0.89). Likewise, 2284 UL specific peaks localized to UL regulatory domains (binomial *P-value*: 5.06×10^−16^), while 1635 localized to DL regulatory domains (binomial *P-value*: 1) (Fig. 5C). In each case, the binomial *P*-value was lower for correlated pairs than for anti-correlated pairs, suggesting that subclass specific peaks are enriched next to subclass specific genes.

### Correlated TFs include known regulators of cell fate in the cortex

Correlated gene expression and chromatin accessibility in a specific subclass may indicate a subclass-specific function. A TF that is highly differentially expressed *and* particularly densely regulated in one subclass for instance may point to a subclass-specific transcriptional network. We therefore searched for densely regulated TFs expressed in DL or UL cells. Using our sets of subclass-specific genes and subclass-specific ATACseq peaks described above, we calculated two binomial *P-values* for each gene: a binomial *P-value* for DL-specific ATACseq peaks and a binomial *P-value* for UL-specific ATACseq peaks. Next, we ranked genes by the log-fold change in the two binomial *P-value*s (Fig. 5E). When we limited the ranking to genes with a log-fold change in gene expression greater than 1 or less than −1, the most densely regulated DL TFs included genes known to be involved in DL cell fate specification (Fig. 5E, ***Supplemental Table 8***) (*Fezf2, NeuroD*, and *Nhlh2*) and enriched in the cortical plate of E13.5 mouse embryos (Figure 6, ***Nhlh1 and Nhlh2***)^57^. Interestingly, the DL TF with the greatest log-fold change in ATACseq *P*-value is the imprinted gene Peg3^58^, although its expression in DL neurons is less than 1-fold higher than in UL neurons.

**Figure 6.**
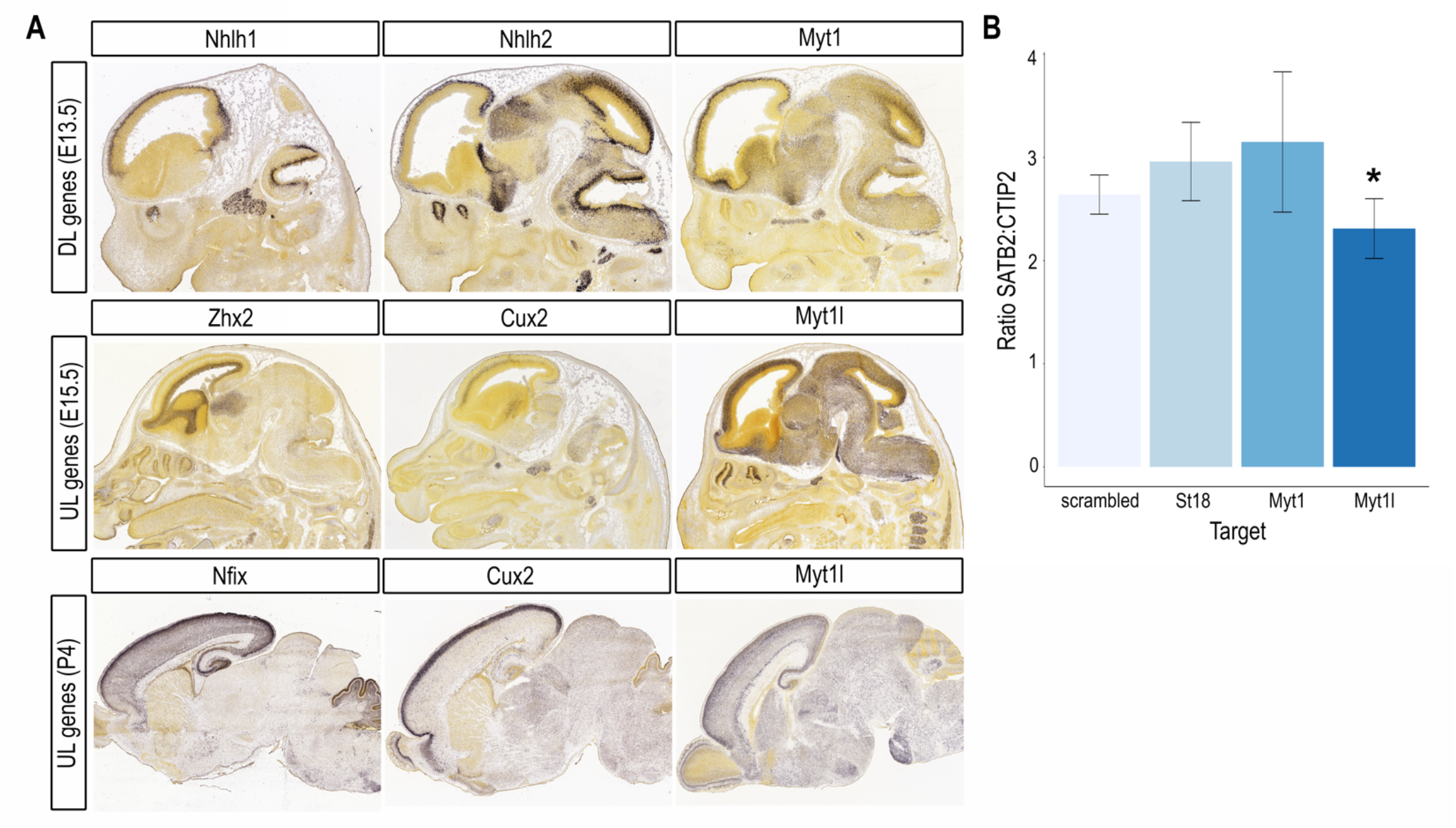
Highly correlated and Myt family TFs show subclass-specific expression in vivo, and Myt1l regulates UL fate in vitro. **(A)** The Myt family TF *Myt1* shows early expression in the mouse cortical plate similar to that of the DL TFs *Nhlh1* and *Nhlh2*, while the Myt family TF *Myt1l* shows early expression in the mouse cortical plate similar to that of the UL TFs *Zhx2* and *Cux2* and enriched UL expression at P4 similar to the UL enrichment of *Nfix* and *Cux2*. *Image credit: Allen Institute* **(B)** Short hairpin RNA (shRNA) knockdown of *Myt1l* but not of *St18* or *Myt1* significantly reduces the ratio of SATB2+ to CTIP2+ cortical neurons.

The most densely regulated UL TFs included genes known to be involved in neural progenitor cell cycle regulation (*Tcf3*^59^, *Tcf4*, and *Hes5*^60^), neuron differentiation (*Nfix* and *Cux2*), or positioning of UL neurons within the cortical plate (*Prdm16*^61^) (***Supplemental Table 8***). We confirmed cortical expression of *Zhx2*, a gene with similar log-fold change in expression and binomial *P-*value of *Cux2*, a major regulator of UL fate (Fig. 5E, Fig. 6A). While *Cux2* expression is enriched in migrating interneurons in the ventricular/subventricular zones, it is also expressed in a subset of radial glial cells in the intermediate zone at E15.5, during UL neurogenesis (Fig. 6A)^62,63^. Similarly, *Zhx2* expression appears enriched in the ventricular zone at E15.5 (Fig. 6A). While UL neurogenesis peaks at E15.5, most UL neurons are not established in the cortical plate until a few days after they exit the cell cycle. We therefore verified UL expression of *Nfix*, the TF with the highest log-fold change in binomial *P-value*, at P4 (Fig. 6A). *Nfix* and *Cux2* are both enriched in UL neurons at P4, suggesting that dense regulation in a specific subclass may indicate a role in cell fate determination.

### Correlated TFs identify subclass-specific gene regulation

To identify potential upstream regulators of densely regulated genes, we searched for enriched TF binding motifs in ATACseq peaks correlated with gene expression in early UL or DL cells. We limited the analysis to genes with a log-fold change in binomial *P*-value greater than 2 and ran seven different published motif discovery tools on each set of correlated peaks^53^. Near identical motif predictions from at least two different tools were combined, and Tomtom^64^ was used to match the most frequently occurring motif with a set of known TF binding motifs.

In each subclass, the top predicted motifs matched TFs found in our set of densely regulated genes (Fig. 5D, E). The top predicted DL motif resembled the consensus motifs for the bHLH TFs NEUROD and NHLH, both of which are more highly expressed and densely regulated in DL neurons. This motif also resembled the TGIF consensus motif, and *Tgif2* is expressed in the ventricular zone of the developing neocortex^65^, with higher expression and chromatin accessibility in early UL neurons (Fig 5E). The top predicted UL motif resembled the consensus motifs for the progenitor TF TEAD2^65,66^, the intermediate progenitor retinoic acid cofactor ZFP423^67^, and the consensus motif for the bHLH oligodendrocyte specification TF OLIG2^68^, all three of which are more highly expressed in early UL neurons (Fig. 5E).

### Myt1l regulates the ratio of upper layer to deep layer neurons

The enrichment of the NHLH binding motif in putative regulatory elements of DL genes suggests involvement of *Nhlh1* and/or *Nhlh2* in DL cell fate. A previous report, however, suggests that both *Nhlh1* and *Nhlh2* are dispensable for normal cortical development^69^. Although this report did not examine expression of layer-specific markers in *Nhlh1/Nhlh2* double-knockout mice, we decided to focus our validation experiments on a family of TFs previously unexplored in the context of cortical cell fate. The myelin transcription factor (Myt) family of zinc-finger proteins includes MYT1/NZF2, MYT1L/NZF1, and ST18/MYT3/NZF3. *Myt1* and *Myt1l* are known to play crucial roles in promoting neuronal differentiation in the telencephalon through repression of neural progenitor genes, thus antagonizing progenitor proliferation^70,71^. Layer-specific function in post-mitotic neurons, however, has not been examined. To determine whether MYT family TFs play a role in cortical cell fate specification, we first confirmed *in vivo* expression of *Myt1* and *Myt1l* in the cortical plate (CP) (Fig. 6A). Similar to *St18* (Fig. 3E), *Myt1* is enriched in the CP at E13.5, when it consists solely of DL neurons. *Myt1l* is enriched in the CP at E15.5 and P4, suggesting expression in both UL and DL neurons. Moreover, *Myt1l* expression clusters with genes that are consistently higher in UL neurons compared with DL neurons in a previous study^21^ (cluster 15).

We tested the function of all three MYT family TFs by shRNA knockdown in cortical neurons cultured from E15.5 mice using previously validated shRNA constructs^71^. To evaluate a difference in the relative abundances of CPN and CFPN neurons, we counted the ratio of SATB2 to CTIP2 four days after transfection (Fig. 6B). Compared with cells transfected with scrambled shRNA-GFP, cells transfected with shRNA to *St18* or *Myt1* showed no significant difference in the ratio of SATB2 to CTIP2. Cells transfected with shRNA to *Myt1l*, however, showed a significant decrease in the ratio of SATB2 to CTIP2 (*P*=0.01), indicating a potential role in promoting UL cell fate.

## Discussion

We have identified distinct transcriptional states of genetically defined deep layer (DL) and upper layer (UL) cortical projection neurons over time. The heterogeneity of each of these bulk populations allowed us to compare intra-group versus inter-group similarity over several stages of development, which revealed that gene expression alone is able to distinguish DL from UL neurons even when limited to transcription factor (TF) genes. Many of these subclass-specific TFs in the mouse likely have conserved function in human brain development given their highly differential expression between embryonic and adult human motor cortex. Moreover, subclass, rather than developmental stage, contributed the highest source of variability in gene expression among DL and UL neurons, and subclass and stage-specific gene expression was significantly correlated with that of genes that define similar populations of sequenced single cells. Therefore, despite the intrinsic heterogeneity of bulk-sorted cells, neurons within each subclass retained more intra-class than inter-class similarity. This phenomenon has been reported as well for subtypes of mature oligodendrocytes that share functional and morphological properties^72^.

The observation that two samples of DL neurons clustered more closely with UL neurons after limiting the analysis to the top 500 most variable transcripts could suggest that DL neurons are more plastic early in development, consistent with the observation that young DL neurons transplanted into an older cortex have the capacity to become UL neurons, while UL neurons transplanted into a younger cortex remain UL neurons^73–75^.

We also identified regions of chromatin that were accessible specifically in one population of cells and correlated with differential gene expression at a given developmental stage. These regions could provide a resource for identifying regulatory elements harboring disease-associated polymorphisms or for understanding the biological role of known polymorphisms associated with neurobiological developmental disorders. Moreover, our finding that subclass-specific accessible chromatin is enriched for subclass-specific TF binding motifs suggests that such polymorphisms could potentially disrupt the function of major transcriptional regulators of cell fate specification and development, and that such erosion of gene regulation could contribute to the mutation load underlying inherited neurological diseases^76^.

Very few regions of accessible chromatin were specific to one population of cells at all three timepoints examined. This finding is perhaps unsurprising given the heterogeneity of the two populations compared. In fact, that hundreds of regions were specific to a subclass throughout several developmental stages is unexpected when considering the vast diversity of cell types in the brain with distinct transcriptional programs^1,77^, and that the majority of open chromatin occupies promoter regions, which often remain open in cell types in which the downstream gene is not expressed. The *St18* promoter, for instance, remains accessible in UL neurons and in mid and late DL neurons even though *St18* is only expressed in early DL neurons (data not shown).

That UL and DL neurons share common expression dynamics reflects the shared biological processes of these two closely related cell types. Conversely, different expression dynamics may indicate cell type-specific functions. Both UL and DL neurons, for example, express long-noncoding RNAs adjacent to major TFs regulating cortical development, suggesting a role for these lncRNAs in cell fate specification, as demonstrated in previous studies^50,78^. *Satb2-lncRNA* expression spikes shortly after neurons reach the cortical plate and start extending their axons from both deep and upper layers. *Satb2-lncRNA* transcripts, however, are differentially localized, confined to the nucleus of SATB2-positive DL neurons but expressed in both the nucleus and cytoplasm of SATB2-positive UL neurons. Moreover, knockdown of *Satb2-lncRNA* caused a subtle but statistically insignificant shift in the ratio of SATB2:CTIP2-postive neurons. More evidence will be required to ascertain whether TF-lncRNAs may play a cell type-specific role in cortical development.

Likely players in the transcriptional networks regulating cell fate are the major TFs adjacent to these TF-lncRNAs, here including *Pou3f2*, *Pou3f3*, and *Satb2*^50^. Numerous studies have suggested that lncRNAs can influence the expression of adjacent genes directly, through recruiting transcriptional or splicing machinery to the locus^78–80^ or binding to chromatin to modulate enhancer-promoter contacts^81^. Alternatively, lncRNA expression can regulate nearby genes indirectly, the process of transcription itself altering chromatin structure in a way that affects expression of the adjacent gene independently of the lncRNA transcript^49, 82–84^. Knockdown of the lncRNA at the mRNA level, therefore, would not affect expression of the nearby regulated gene. The small but not significant reduction in the number of SATB2-positive cells relative to CTIP2-positive cells after siRNA knockdown of *Satb2-lncRNA* could indicate that *Satb2-lncRNA* is only a byproduct of transcription through the *Satb2* locus, that other players in the transcriptional network compensate for the reduction in *Satb2-lncRNA*, that *Satb2* splicing or transport is affected in a way that does not change total SATB2 levels, or that residual *Satb2-lncRNA* in the nucleus is enough for the system to function normally.

Additional players in the transcriptional networks regulating cell fate in the developing cortex could include TF families previously unexplored in this context. One such family is the Myt family of TFs, shown here to have dynamic expression patterns throughout cortical development: *St18* expression in early DL mouse neurons reflects its expression in the developing human motor cortex, while *Myt1* and *Myt1l* expression in DL versus UL neurons suggests opposing roles in cell fate specification. Indeed, we found that knockdown of *Myt1l* in cortical neurons caused a significant reduction in the ratio of SATB2:CTIP2-positive cells. Moreover, one isoform of *Myt1l* (*ENSMUST00000092649*) is significantly higher in UL neurons at all three stages examined here. Compared to full length *Myt1l*, this truncated isoform lacks several SIN3 interaction domains, which are important for repression, but retains most of its zinc finger domains, suggesting that its role in cortical neuron development may differ from its characterized role in repressing non-neural fates in progenitor cells.

Here we have generated a resource for discovering genes and regulatory elements important for establishing subclasses of cells in the developing mouse cortex. The recent finding that genes involved in chromatin changes during learning and memory are known to be involved in brain development and associated with autism spectrum disorders^85^ suggests that developmental regulatory elements harboring disease-associated polymorphisms may play a dual role in synaptic plasticity. Identifying and understanding the role of such regulatory elements during development could therefore improve our ability to characterize their roles in disease pathogenesis during adulthood.

## Acknowledgments

We thank Dr. Chris Kaznowski for technical help, Dr. Moritz Mall for generously sharing shRNA constructs, Dr. Paola Arlotta for the Cdk5r-GFP construct, Dr. Ben Barres for insights regarding OPCs, and members of the McConnell and Bejerano labs for helpful comments, especially Dino Leone and Pushkar Joshi. This work was supported by a National Institute of Neurological Disorders and Stroke (NINDS) Epilepsy Training Grant (W.E.H.), a Stanford Neurosciences Institute Interdisciplinary Scholar Postdoctoral Fellowship (W.E.H.), a National Institutes of Health (NIH) R01 MH51864 (S.K.M.), and a Stanford Bio-X Interdisciplinary Seed grant (G.B. and S.K.M.).

## Materials and Methods

### FACS purification of cortical neuron sub-populations

All animal work was carried out in compliance with Stanford University IACUC under approved protocol #11499 and institutional and federal guidelines. The day of vaginal plug detection was designated as E0.5. The day of birth was designated as P0. For deep layer neurons, CD1 female mice were crossed with homozygous golli-τ-EGFP males, resulting in heterozygous offspring. For upper layer neurons, a plasmid encoding Cdk5r-promoter-GFP (***generous gift of Dr. Paola Arlotta***) was introduced into the developing neocortex of embryos of timed-pregnant CD1 dams on E15.5 (4-7 embryos per litter were injected) by *in utero* electroporation as previously described^86^.

For each biological replicate, the neocortex of one litter of embryos (two to five per litter) was microdissected in cold HBSS. Tissue was dissociated into a single-cell suspension using papain according to the manufacturer’s instructions (Worthington), and resuspended in cortico-spinal motor neuron (CSMN) medium (1mM pyruvate, 2mM L-glutamine, 5ug/mL insulin, 100U/100ug/ml pen/strep, 1X Sato, 35mM glucose, 0.34% BSA, and 800 uM kynurenic acid in 50% DMEM/50% Neurobasal)^87^. FACS purifications were performed on a BD FACSAria II, BD FACSJazz, BD Influx, or BD FACS Aria Fusion. Gates for FACS were set using non-GFP age-matched or littermate controls and confirmed using immunohistochemistry for GFP on pre- and post-sorted cells. For RNA isolation, 75,000-500,000 cells were used per biological replicate. For accessible chromatin, 25,000-50,000 cells were used per biological replicate. Sorted cells were stored in cold CSMN and immediately processed for RNA or accessible chromatin isolation.

### Immunohistochemistry and Fluorescent In Situ Hybridization

Immunohistochemistry was performed as previously described^18^. Briefly, whole embryos or brains from animals older than E17 were dissected in cold PBS and fixed overnight in 4% paraformaldehyde in 0.1 M phosphate buffer pH 7.3, cryoprotected in 30% sucrose, and frozen in OCT. Frozen tissue was cryosectioned at 14 µm on a Leica CM1900 cryostat, and sections were incubated overnight at 4°C with the following primary antibodies: rabbit anti-GFP (Invitrogen A-11122, 1:1000), mouse anti-Satb2 (Abcam ab51502, 1:200), and rat anti-Ctip2 (Abcam ab184675-100, 1:300). Alexa Fluor-conjugated secondary antibodies, including goat anti-rat 647, goat anti-mouse 555, and goat anti-rabbit 488, were used to detect primary antibodies. Fluorescent *in situ* hybridization was performed using the RNAscope Fluorescent Multiplex Assay and probes to *St18*, *Satb2*, and *Satb2-lncRNA* according the manufacturer’s instructions (ACDBio). Images were taken on a Nikon 80i microscope with a Hamamatsu Orca ER camera or a Zeiss LSM 510 meta confocal microscope and postprocessed using Adobe Photoshop CS3.

### RNAseq and ATACseq Library Preparation and Sequencing

Bulk RNA was isolated from sorted cells using an RNAeasy Micro Kit (Qiagen) according to the manufacturer’s instructions. RNA quality was assessed using a Bioanalyzer 2100 (Agilent), and the RNA integrity number for each sample was above 9. At least 100ng of total RNA input was used per library. Libraries for mRNA sequencing were prepared using a TruSeq Stranded mRNA Sample Prep Kit (Illumina), and libraries were sequenced on a HiSeq2500 (Illumina), generating an average of 2.35×10^7^ 100bp paired-end reads per library (range, 1.65×10^7^ − 2.86×10^7^). Accessible chromatin was isolated as described in *Buenrostro et al.* (2013), with the following modification: twelve rounds of PCR were used for all samples. Resulting ATACseq libraries were sequenced on a HiSeq2500 (Illumina), generating an average of 1.03×10^8^ 100bp paired-end reads per library (range 2.51×10^7^ – 1.91×10^8^).

### RNAseq Analysis

Paired-end reads were aligned to the mouse genome (mm9) using kallisto version 0.43.1^88^ with default parameters and a transcriptome index built using Ensembl transcripts. An average of 2.35×10^7^ reads were uniquely mapped (range, 1.65×10^7^ – 2.86×10^7^). Transcript abundance and differential expression were determined using Sleuth^89^.

For lncRNA analysis, paired-end reads were aligned to the mouse genome (mm9) using STAR version 2.5.3^90^ with default parameters for an average of 2.35×10^7^ uniquely mapped reads (range, 1.70×10^7^ – 2.99×10^7^). Transcripts were assembled using Cufflinks v2.2.1^91^ and gene models built using UCSC transcripts masked for rRNAs, tRNAs, and mitochondrial genes. Transcript abundance and differential expression were determined using Cuffdiff 2^92^.

### ATACseq Analysis

Reads were trimmed using cutadapt version 1.9^93^ and pairs were aligned to the mouse genome (mm9) using Bowtie2 version 2.2.6^94^ for an average of 1.01×10^8^ reads per biological replicate (range, 2.46×10^7^ – 1.86×10^8^). Duplicate reads were removed using the Picard Tools version 1.140 MarkDuplicates function for an average of 7.11×10^7^ reads per biological replicate (range, 1.32×10^7^ −1.56×10^8^). Peaks were called using MACS version 2.1.0 with a *P*-value cutoff of 0.01. Replicated peaks were defined as overlapping regions within overlapping peaks and merged using bedtools. Peaks were masked for blacklist regions, segmental duplications, repeat regions, and exons. ATACseq peaks were associated with genes using basal regulatory domains defined by GREAT (version 2.0.2) as 5 kb upstream and 1 kb downstream plus up to one Mb in both directions to the nearest gene’s basal regulatory domain^55^.

### Hierarchical Clustering, PCA and tSNE Analysis

Naïve hierarchical clustering was performed using the heatmap.2 function in R. For PCA, starting with a transcriptome of 95883 Ensembl transcripts, the TPM of all transcripts for a given gene were summed. This reduced each RNA-seq quantification to a vector of 34307 values, corresponding to 34307 genes. Across both replicates for all six conditions (3 UL and 3 DL), this yielded 12 x 34307-vectors for PCA, which was conducted using sciKitlearn^95^ version 0.18.1. For PCA on transcription factors only, we repeated this process, but only considered the 1461 genes corresponding to our transcription factor library^96^.

TPM measurements of expression over Ensembl transcripts at each time point were encoded in a 95883 dimensional vector. First, a PCA reduction was performed to reduce the representation to twelve dimensions then the t distributed stochastic neighbor embedding (t-SNE) algorithm^97^ with perplexity .7 was used to dimensionally reduce the TPM vectors to two dimensions.

### Gene Ontology Analysis

Shared biological function of genes that change in the same direction in both cell subclasses was determined using the Gene Ontology Consortium Database on 20 February 2018^45,98^. Molecules annotated for axon guidance were identified using the Mouse Genome Database at the Mouse Genome Informatics website, The Jackson Laboratory, Bar Harbor, on 28 September 2017.

### Evolutionary Conservation and Motif Discovery

To calculate top conserved peaks in upper layers, we first split p300 peaks from E14.5 neocortex into windows of 50 base pairs and annotated each window with the sum of PhastCons^99^ scores for each base pair. Each base pair is assigned a PhastCons score between 0 and 1. Thus, each 50-basepair-window can receive a maximum PhastCons score of 50. All 50-base pair-windows with a summed PhastCons score greater than 40 were retained. Subsequently, all retained adjacent 50-basepair windows (with a maximum gap of 20 base pairs between adjacent windows) were merged into larger, highly conserved, genomic regions. We refer to this set of genomic regions as “highly conserved p300 regions”.

Next, ATACseq peaks from both replicates of E13 DL and E17 UL were intersected to arrive at a set of DL peaks and a set of UL peaks. Subsequently, all highly conserved p300 regions were selected based on overlap with DL or UL peaks. Only highly conserved p300 regions overlapping DL or UL peaks were retained to arrive at a set of “highly conserved p300 regions overlapping ATACseq peaks; all highly conserved p300 regions that did not overlap at least one intersected peak were discarded. Further, all highly conserved p300 regions overlapping ATACseq peaks were associated with the nearest genes by distance from the gene’s transcription start site (TSS). All regions overlapping a TSS (distance from TSS equals zero) were discarded to arrive at a set of “conserved DL and UL p300 enhancers”. UL-specific enhancers were defined as conserved UL p300 enhancers not overlapping conserved DL p300 enhancers, and vice versa. Both sets of enhancers were sorted by the summed PhastCons score divided by distance from the nearest gene’s TSS (descending).

TF binding site prediction was performed as described previously^53^. Briefly, we ran seven different motif discovery tools on each set of correlated peaks, using the set of correlated peaks in the comparison cell subclass as the background set. Tools included AlignAce^100^, CisFinder^101^, MDscan^102^, MEME^103^, MoAn^104^, MotifSampler^105^, and Weeder^106^. Near identical motif predictions from at least two different tools were combined and compared with a database of known TF binding motifs using TomTom version 5.0.0^64^.

### Primary cell culture, siRNA knockdown of lncRNA, and transfection of lentiviral constructs

For siRNA knockdown of Satb2-lncRNA, the neocortex was dissected on embryonic day 15, cells were dissociated using trypsin, cultured in NeuroCult basal medium containing EGF and proliferation supplement (Stemcell 05702) overnight, and switched to differentiation medium (Stemcell 05704) on the following day. Four hours after switching to differentiation medium, cells were transfected using Lipofectamine RNAiMax reagent (Invitrogen) according to the manufacturer’s instructions. After 4 days in culture, cells were fixed with cold 4% PFA for 5 minutes then processed for IHC. The siRNA sequence for “siRNA 1” was ACTCACTGACAAGCCGCAGAGAGAA, for siRNA 2 was GAGATGATTATTAGTTGCGTTGAGT, for scrambled 1 was ACTTCAGACCCGGAGAAACGAA, and for scrambled 2 was GAGTTAGATTAGTTGTTGCGTAAGT (Stealth RNAi, ThermoFisher). Knockdown was confirmed using qPCR and primers to Satb2-lncRNA (tgatcaAGACCGGTTCTGGAGAGAAAG) short isoform (ctcgaTGAATCATCATCAAATATATTTATTCACTG) or long isoform (ctcgagTATATATGTTTAATTACACAGTAGTAGAACAT).

For shRNA knockdown of Myt family TFs, dissections and infections were performed as described above with the following changes: shRNA directed to *St18*, *Myt1*, or *Myt1l* (***Supplemental Table 9***) cloned into a pSico-GFP lentiviral vector (generous gifts of Dr. Moritz Mall) was introduced to cultured cells four hours after switching to differentiation medium. Two different sequences were used for each target. After 4 days, cells were fixed with cold 4% PFA for 5 minutes and processed for IHC.

For cell counts, a minimum of 3 biological replicates per construct were counted, and a minimum of 100 cells per image was required for inclusion in the final analysis. A Student’s T-test was used to determine significance.

---

**Figure S1.**
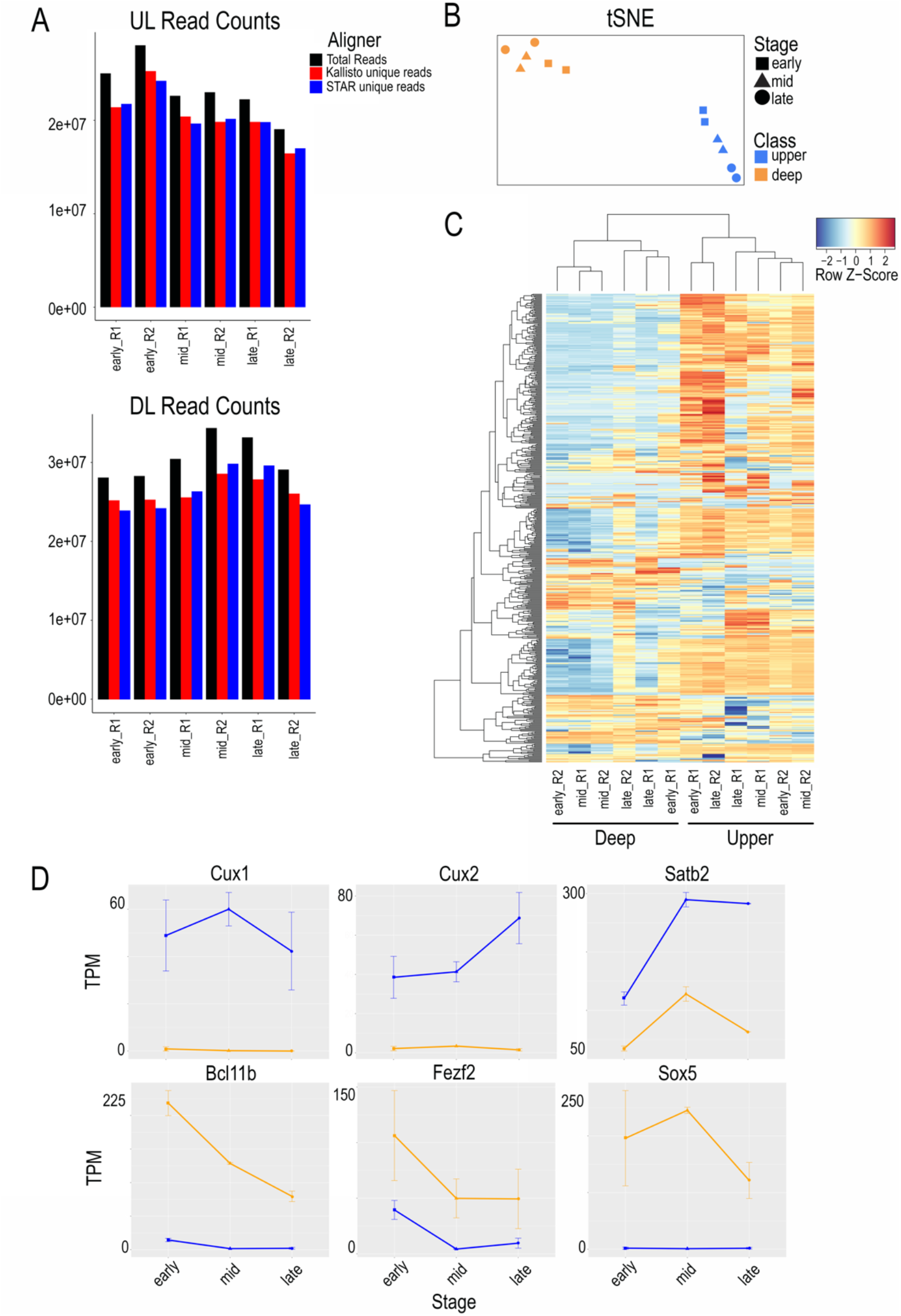
Cell type is the greatest source of variability in gene expression. **(A)** Total and aligned read counts for each RNAseq library are similar for UL and DL populations. **(B)** t-distributed stochastic neighbor embedding (t-SNE) of all transcripts shows segregation of UL and DL populations. **(C)** Naive hierarchical clustering of the top 500 most variable transcription factor transcripts shows segregation of UL and DL populations. **(D)** Transcripts per million (TPM) of select TFs known to be more highly expressed in either UL neurons (*Cux1*, *Cux2*, and *Satb2*) or DL neurons (*Bcl11b*, *Fezf2*, and *Sox5*) are highest in the appropriate cell class for each TF.

**Figure S2.**
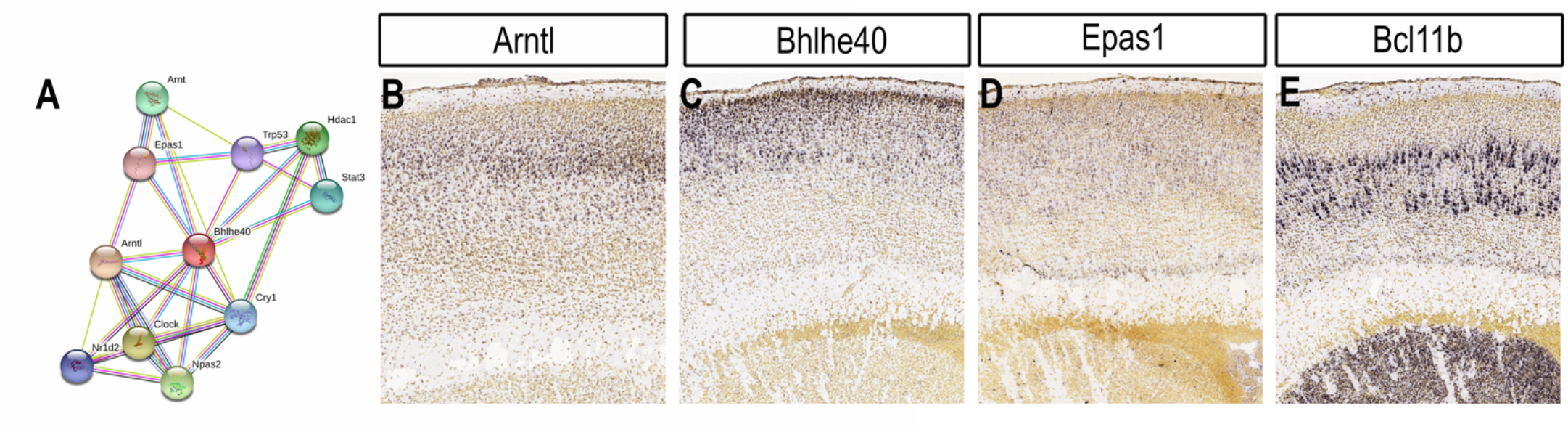
A subnetwork of circadian genes is expressed in developing UL neurons. **(A)** Predicted protein-protein interactions of the UL TF BHLHE40. *Source: STRING database*. **(B-D)** *In situ* hybridization of three circadian-associated genes showing enriched expression in UL neurons of the mouse neocortex at P4. *Image credit: Allen Institute*. **(E)** Layer 5 expression of *Bcl11b* at P4 showing exclusion from UL cells. *Image credit: Allen Institute*.

**Figure S3.**
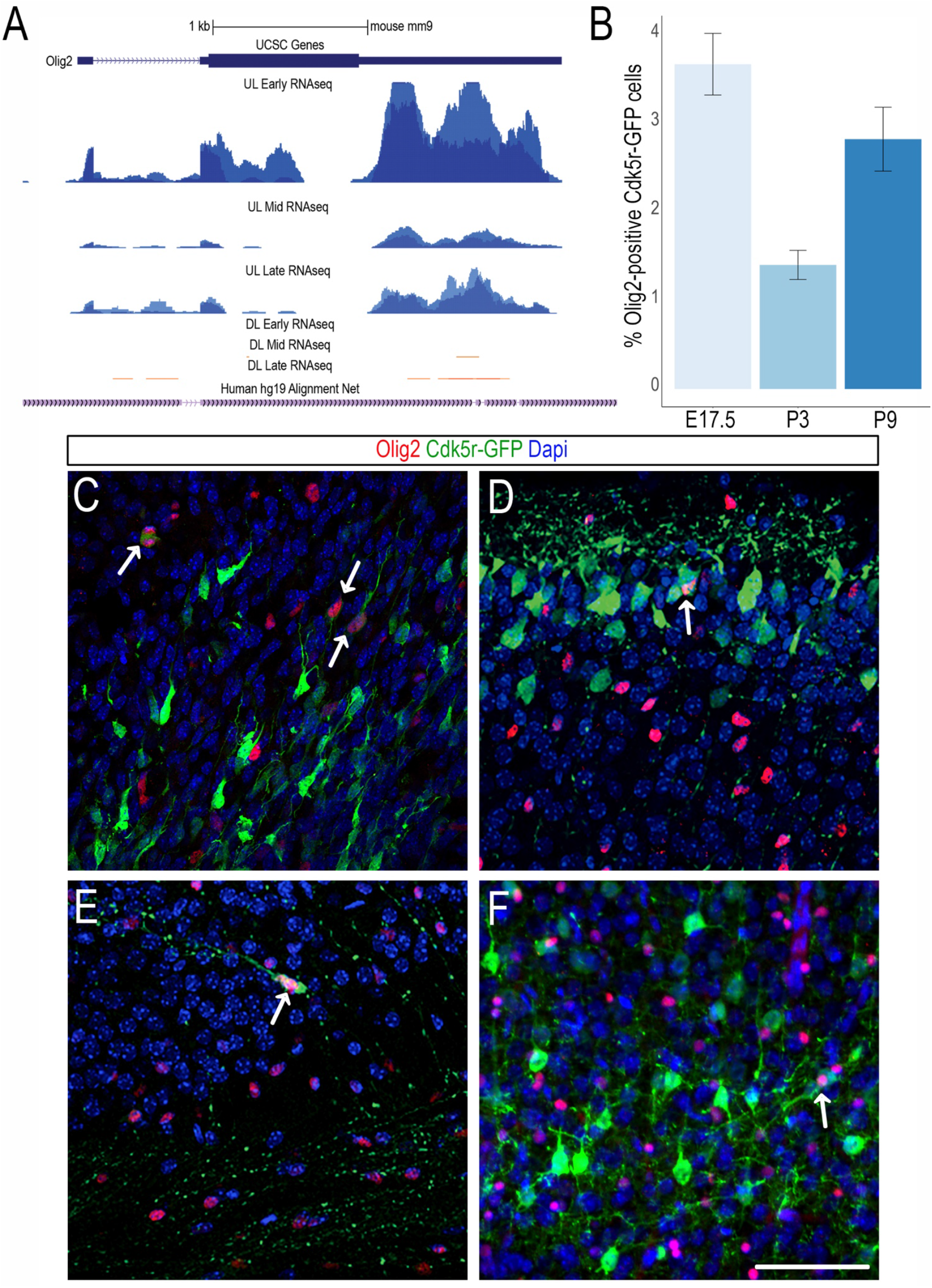
A subset of Cdk5r-positive cells expresses Olig2. **(A)** RNAseq read pileups show highest expression of *Olig2* RNA in early Cdk5r-GFP+ compared with mid and late, when reads appear concentrated in the 3’ UTR. **(B)** The percentage of Olig2+ Cdk5r-GFP+ cells is between 1% and 4% at all three stages. **(C-F)** Immunohistochemistry for OLIG2 shows expression in a subset of Cdk5r-GFP+ cells (arrows) in E17.5 mouse neocortex (C), P3 mouse neocortex (D), P3 mouse cingulate cortex (E), and P9 mouse neocortex (F). Scale bar in F is 50 μm.

**Figure S4.**
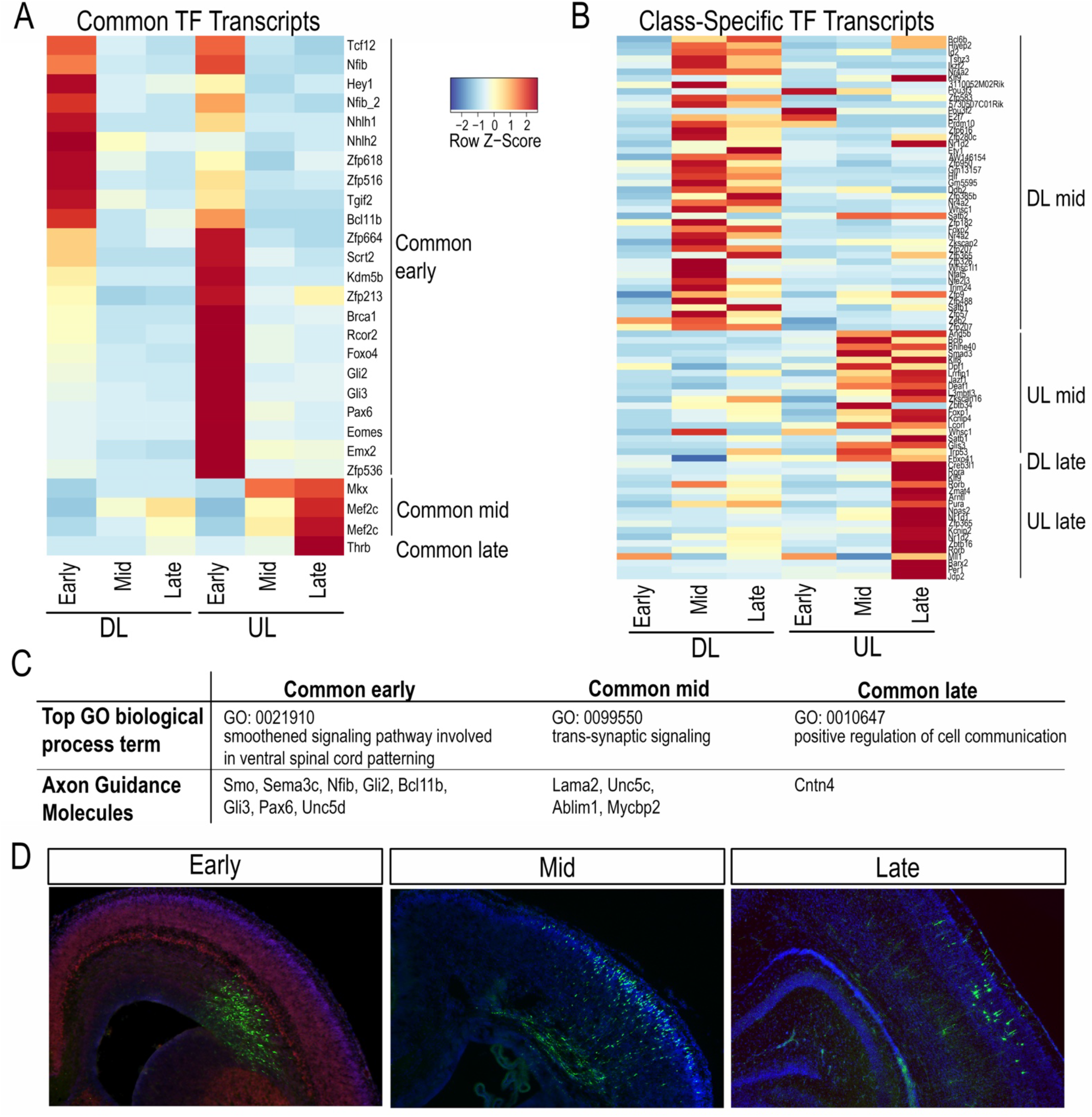
DL and UL neurons have common and specific expression dynamics. **(A)** A subset of TFs is expressed early, mid, or late in both DL and UL neurons. **(B)** A subset of TFs is expressed mid or late only in DL or UL neurons. **(C, D)** The top gene ontology term enriched for all common early, mid, and late transcripts reflects known stage-specific biological processes, and select axon guidance molecules are expressed at each stage in both subclasses (C). Mouse cortical neurons electroplated with Cdk5r-GFP (green) at E15.5 show morphological changes from E17.5 (early), before they enter the cortical plate marked by MYT1L (red), to P1 (mid), during primary axon extension, and P5 (late), during axon collateral formation.

**Figure S5.**
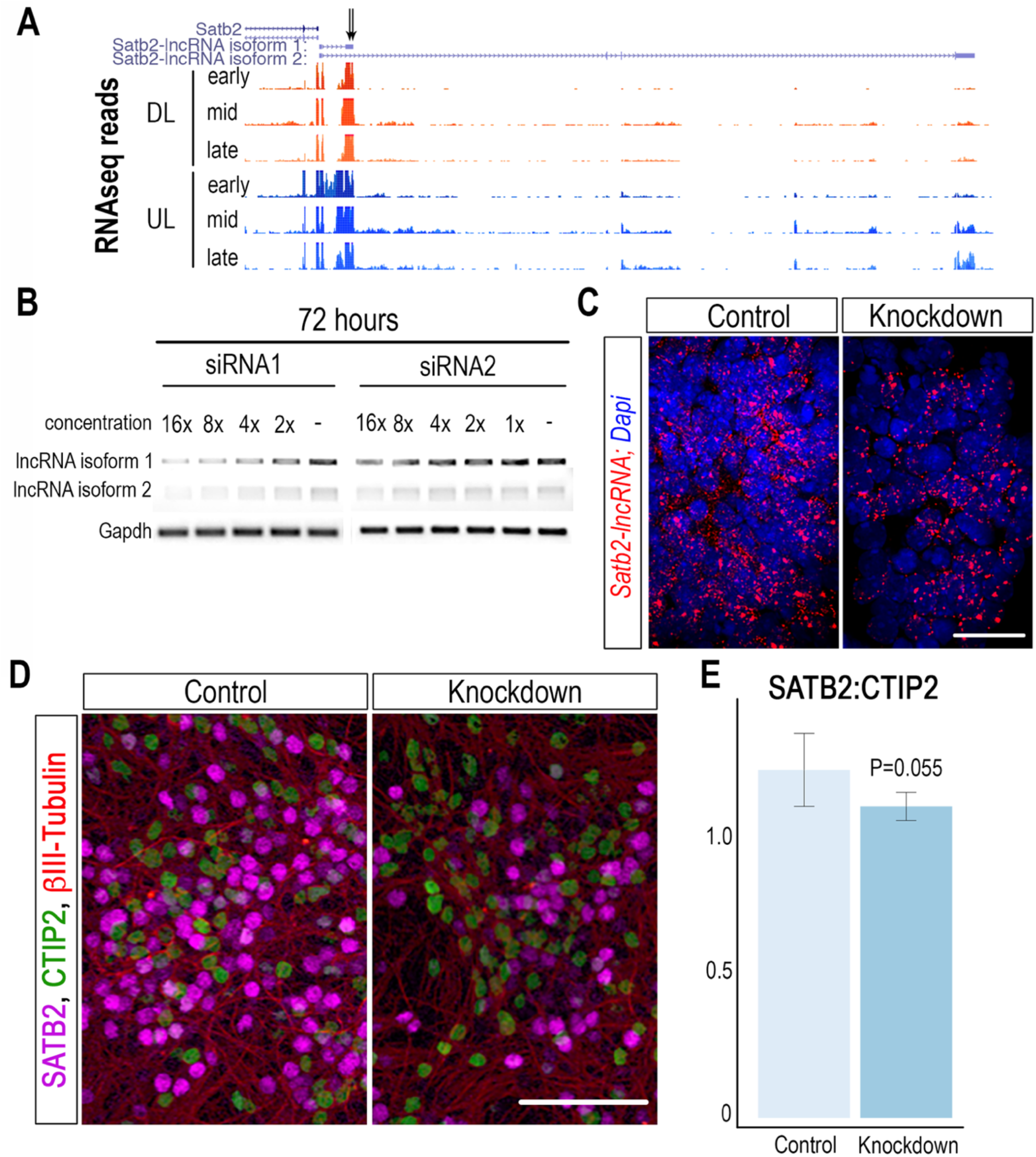
Cytoplasmic knockdown of two isoforms of *Satb2-lncRNA* in cortical neurons. **(A)** RNAseq read pile-ups show expression of the short and long isoforms of *Satb2-lncRNA* in cortical neurons. **(B)** Two different siRNA constructs targeted to the first exon of *Satb2-lncRNA* reduce expression of both the short and long isoform at 16X concentration. **(C)** Fluorescent *in situ* hybridization of *Satb2-lncRNA* (red) shows reduced cytoplasmic expression after siRNA treatment. **(D-E)** Dissociated cortical cells treated with vehicle or siRNA to *Satb2-lncRNA* stained with SATB2, CTIP2, and β−III-Tubulin show a reduced ratio of SATB2 to CTIP2 that only trends toward significance. Scale bar in C is 25 μm. Scale bar in D is 100 μm.

